# *EROS* is required for phagocyte NADPH oxidase function in humans and its deficiency causes Chronic Granulomatous Disease

**DOI:** 10.1101/331454

**Authors:** David C. Thomas, Louis-Marie Charbonnier, Andrea Schejtman, Hasan Aldhekri, Eve Coomber, Elizabeth R. Dufficy, Anne E. Beenken, James C. Lee, Simon Clare, Annaliese O. Speak, Adrian J. Thrasher, Giorgia Santilli, Hamoud Al-Mousa, Fowzan S. Alkuraya, Talal A. Chatila, Kenneth G.C. Smith

**Affiliations:** Department of Medicine, University of Cambridge School of Clinical Medicine, Box 157 Cambridge Biomedical Campus, Cambridge, CB2 0QQ, United Kingdom; Division of Immunology, Boston Children’s Hospital and Department of Pediatrics, Harvard Medical School, Karp Family Building, Room 10-214, 1 Blackfan Street, Boston, MA, 02115, USA; Molecular Immunology Unit, UCL Great Ormond Street Institute of Child Health, 30 Guilford Street, London, WC1N 1EH; Department of Paediatrics, King Faisal Specialist Hospital and Research Center, Riyadh, Saudi Arabia, PO Box 3354, MBC 58; Wellcome Trust Sanger Institute, Wellcome Trust Genome Campus, Hinxton, Cambridgeshire, CB10 1SA, United Kingdom; College of Medicine, Alfaisal University, Riyadh, Saudi Arabia; Department of Genetics, King Faisal Specialist Hospital and Research Centre, Riyadh, Saudi Arabia; Department of Anatomy and Cell Biology, College of Medicine, Alfaisal University, Riyadh, Saudi Arabia; Saudi Human Genome Programme, King Abdulaziz City for Science and Technology, Riyadh 12371, Saudi Arabia

**Keywords:** EROS, C17ORF62, CYBC1, Chronic granulomatous disease, Nox2, gp91*phox*

## Abstract

The phagocyte respiratory burst is mediated by the phagocyte NADPH oxidase, a multi-protein subunit complex that facilitates production of reactive oxygen species and which is essential for host defence. Monogenic deficiency of individual subunits leads to chronic granulomatous disease (CGD), which is characterized by an inability to make reactive oxygen species, leading to severe opportunistic infections and auto-inflammation. However, not all cases of CGD are due to mutations in previously identified subunits. We recently showed that Eros, a novel and highly conserved ER-resident transmembrane protein, is essential for the phagocyte respiratory burst in mice because it is required for expression of gp91*phox*-p22*phox* heterodimer, which are the membrane bound components of the phagocyte NADPH oxidase. We now show that the function of *EROS* is conserved in human cells and describe a case of CGD secondary to a homozygous *EROS* mutation that abolishes EROS protein expression. This work demonstrates the fundamental importance of *EROS* in human immunity and describes a novel cause of CGD.

**Clinical Implications:** Chronic granulomatous disease is caused by an inability to make reactive oxygen species via the phagocyte NADPH oxidase. Mutations in C17ORF62/EROS, which controls gp91*phox*- p22*phox* abundance, are a novel cause of chronic granulomatous disease.

**Key Messages:**

- The murine gene, *Eros*, is known to regulate abundance of gp91*phox*-p22*phox* heterodimer and *Eros* deficient mice are susceptible to infection
- We show that the function of *EROS* is conserved in human cells and that a homozygous mutation in EROS causes chronic granulomatous disease

## Results

The multi-subunit phagocyte NADPH oxidase generates reactive oxygen species and is crucial for host defence^1^. Deficiencies in individual subunits (gp91*phox*, p22*phox*, p47*phox*, p67*phox* and p40*phox*) cause chronic granulomatous disease (CGD) but some patients with CGD do not have mutations in these genes^2^. We recently found that Eros^3^, a hitherto undescribed protein, is essential for the generation of reactive oxygen species because it is necessary for protein (but not mRNA) expression of the gp91*phox*-p22*phox* heterodimer, which is almost absent in *Eros*-deficient mice. *Eros*-/- animals succumb quickly following infection with *Salmonella* Typhimurium or *Listeria Monocytogenes*. *Eros* is highly conserved^3^ and has a human orthologue *C17ORF62 (*hereafter referred to as *EROS)*. We asked whether the gene fulfilled the same function in humans. We performed CRISPR-mediated deletion of *EROS* in PLB-985 cells, (**Fig. S1A**) and identified two clones with 8bp and 1bp deletions respectively (**Fig. S1B and C**). Neither clone expressed EROS protein (**Fig. 1A**) or detectable gp91*phox* (**Fig. 1B**). p22*phox* expression was also much lower in both *EROS*-deficient clones than in control cells (**Fig. 1C**). We verified the lack of surface gp91*phox* expression by flow cytometry (**Fig. 1D**). Both *EROS*-deficient clones had a severely impaired respiratory burst (**Fig. 1E**). In addition, *EROS*-deficient clones differentiated towards a neutrophil phenotype also demonstrated an impaired respiratory burst (**data not shown**). As expected, re-introduction of EROS using a lentiviral vector restored gp91*phox* expression to EROS-deficient clones **(Figure 1F**) and oxidase activity as measured by Nitro blue tetrazolium chloride (NBT) test (**Figure 1G**) and DIOGENES assay (data not shown).

**Figure 1:**
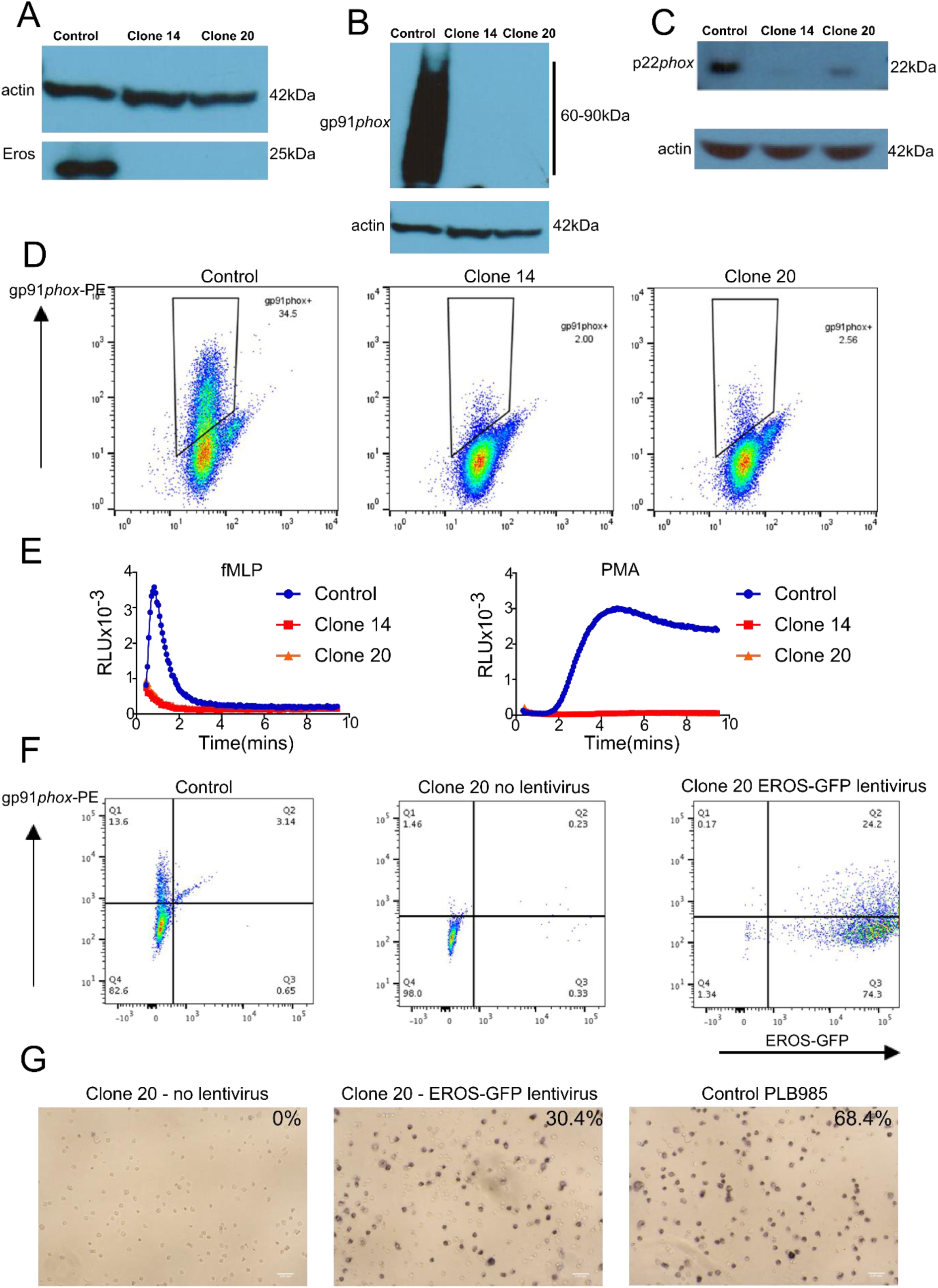
The function of EROS is conserved in humans. (**A-C**) Western blot for expression of (**A**) EROS protein (**B**) gp91*phox* **(C)** p22*phox* in the CRISPR targeted PLB-985 clones. (**D**) flow cytometry for expression of gp91*phox* on the plasma membrane of control and CRISPR targeted PLB-985 clones. **(E)** assessment of the phagocyte respiratory burst by DIOGENES in non-matured PLB-985 cells and **(F)** flow cytometry for expression of gp91*phox* on the plasma membrane of CRISPR targeted PLB-985 with or wthout lentiviral overexpression of EROS-GFP. **(G)** Differentiated WT PLB985, EROS KO (clone 20) and clone 20 previously reconstituted with the EROS-GFP lentiviral vector (86% GFP^+ve^) were incubated with 100ul of 1mg/ml NBT solution containing 1ug/ml of PMA for 30 minutes minutes at 37C. Oxidase positive cells (blue cells) were scored in 4 different fields. Data are representative of 3 independent experiments.

We then identified a patient with a homozygous *C17ORF62/EROS* mutation in a resource paper that details a thousand Saudi Arabian families with genetic disease^4^. He presented with fever, splenomegaly, lymphadenopathy and short stature, but no immuno-phenotyping was detailed at that time^4^. His full clinical history is as follows. He is a Saudi Arabian boy, born in 2007, the son of parents in a consanguineous marriage. He has three healthy older sisters. At 2 months of age, he developed a localized abscess following BCG vaccination. He was then relatively well until 8 years of age but was noted to be of short stature and experienced recurrent pulmonary infections and tonsillitis/pharyngitis despite tonsillectomy.

In August 2015, he became unwell with a febrile illness and an abnormal dihydrorhodamine (DHR) test was noted (**Fig. 2A,B**). He has a severely impaired DHR in response to both PMA and zymosan. He was also profoundly lymphopenic. He subsequently developed an acute episode of hemolytic anemia in November 2015 and again in January 2017. During 2015, his fevers, infection, lymphopenia and elevated inflammatory markers met criteria for a diagnosis of haemophagocytic lymphohistiocytosis (HLH). He had no mutations previously implicated in HLH pathogenesis. He was treated with cyclosporine and steroids and was transferred to Boston Children’s Hospital (BCH) in December 2016 for consideration of bone marrow transplantation. The DHR test was repeated and rhodamine fluoresence was again virtually absent. Further assessment demonstrated granulomatous inflammation in his lungs, and discrete granulomata in his bone marrow, with no evidence of infection. Following his open lung biopsy at BCH, he developed hemolytic anemia and required Intensive Care. He recovered with steroids, however on weaning of this therapy he developed recurrent pleural effusions. Due to his autoimmunity, features of lymphopenia with granulomata, and pleural effusions/hemolytic anemia he was started on sirolimus and he remains in this therapy. His steroids have been weaned to 4mg daily. While reasonably well clinically, he developed diminished anti-pneumococcal antibody responses, worsening lymphopenia, and declining immunoglobulin levels. He therefore underwent a myeloablative bone marrow transplant and has recovered well.

**Figure 2:**
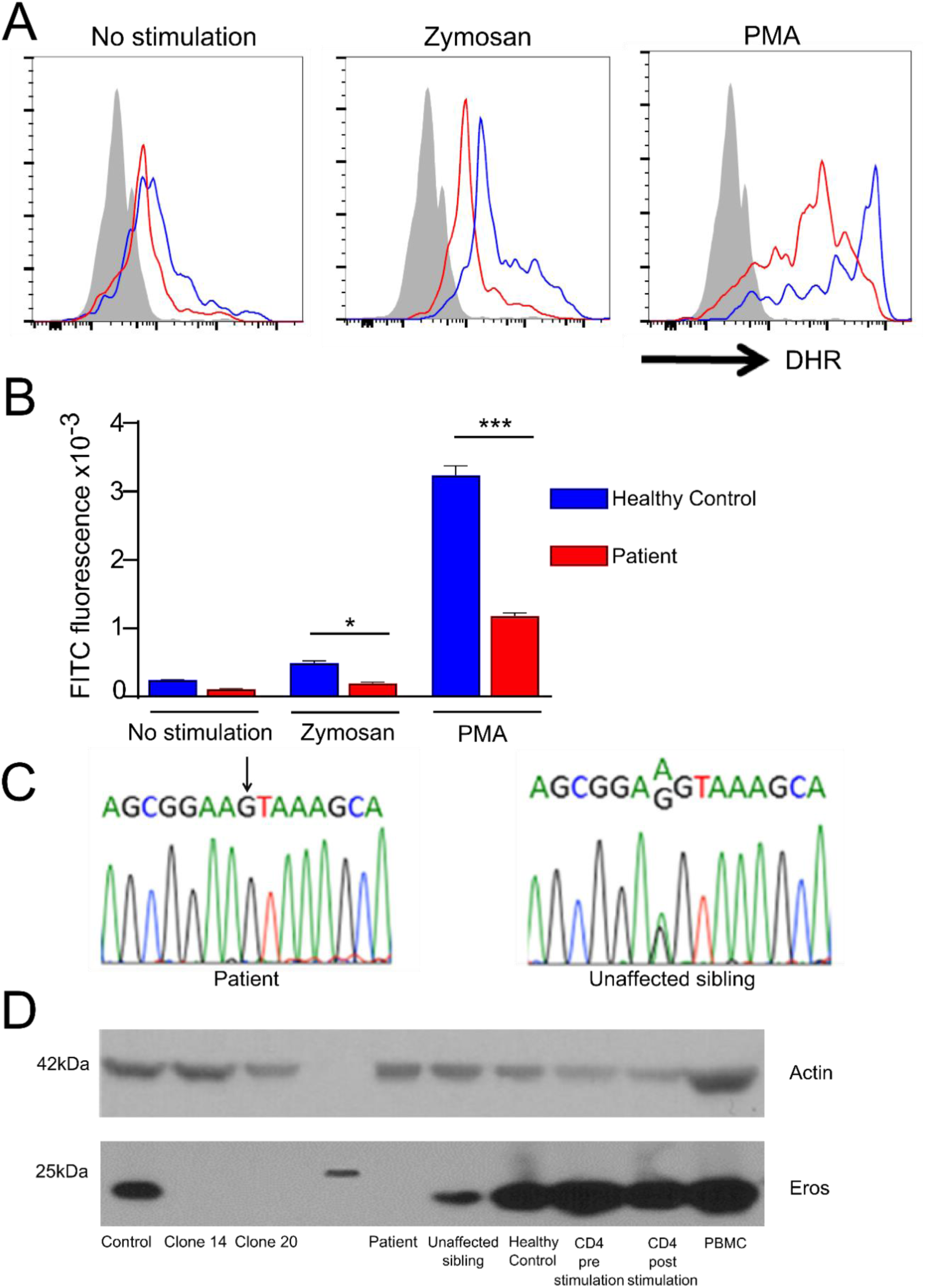
A homozygous *EROS* mutation leads to loss of protein expression and an impaired phagocyte respiratory burst. **A, B)**: phagocyte respiratory burst in response to the indicated stimuli in neutrophils isolated from the patient and a healthy control. **(C)** homozygous c.127 A to G mutation in the patient and the same sequence in his heterozygous sister **(D)** EROS protein expression in PLB-985 cells and CRISPR targeted clones and in CD3/CD2/CD28 expanded PBMC from the patient, his sister and a healthy control. EROS protein expression in primary CD4+ T cells pre and post CD3/CD2/CD28 and PBMC from a further healthy control are also shown in this blot **(C and D).** Data in A, C and D are representative of 2-3 independent experiments.

Whole exome sequencing demonstrated that the patient had a homozygous (c.127 A to G, NM_001033046) mutation in *C17ORF62* (human *EROS*). His sisters were all heterozygous for this mutation, confirmed by Sanger sequencing (**Fig. 2C**). Based on analysis of other family members and the likely important role of *C17ORF62 (EROS)* in immunity, this mutation was identified as the most likely cause of the patient’s disease. The mutation was not present in 10,000 whole genomes from the United Kingdom National Institute for Health Research BioResource - Rare Disease cohort (which includes 1000 patients with primary immunodeficiency), nor in gnomAD. The variant was also absent in 3,300 ethnically matched exomes. It is, therefore, not seen in seen across 110,579 individuals with sequence data coverage across this position. There were no deleterious mutations in known NADPH oxidase subunits.

Splice site prediction algorithms including Mutation Tasting (**www.mutationtaster.org**) and Human Splicing Finder (**http://www.umd.be/HSF3/**) predict that the variant both disrupts an exonic splice enhancer (ESE) and creates an exonic splice silencer (ESS) which is likely to lead to a retained intron. This intron has 4 in-frame stop codons. It is therefore likely that translation would cease downstream of exon 4. Even if splicing is not disrupted, PolyPhen (**http://genetics.bwh.harvard.edu/pph2/**) predicts that the D43N mutation that would occur in the translated protein is also damaging.

We therefore performed western blot analysis on anti-CD3-CD28-CD2 expanded T cells from the patient, his sister and a healthy control, as well as primary T cells (either pre or post polyclonal stimulation) and peripheral blood mononuclear cells from healthy volunteers. The CRISPR targeted clones described above were used as positive and negative controls respectively. The patient had undetectable levels of EROS protein compared with cells from the healthy control or the primary T cells/PBMC, while his heterozygous sister had intermediate levels (**Fig. 2D**).

## Discussion

This work demonstrates that the function of the novel transmembrane protein, Eros, is conserved in humans. It also represents the first description of an immunodeficiency syndrome secondary to mutations in *C17ORF62 (*human *EROS)*. The severity of the disease seen in this patient underlines the importance of human *EROS* is for normal immunity. The patient has a clinical history that is compatible with a diagnosis of autosomal recessive CGD, in that he had both infectious and auto-inflammatory manifestations, together with histopathological evidence of granuloma formation in the context of an impaired DHR response. While recurrent infections, BCG-itis^2,5^, granulomatous inflammation and HLH^6^ are all recognised sequelae of CGD, this patient some unusual features such as autoimmune haemolytic anaemia. This is uncommon in CGD and may represent an effect of *EROS*- deficiency that is independent of its effects on the NAPDH oxidase.

The high sequence similarity between mouse and human *EROS* and the loss of gp91*phox* expression and the phagocyte respiratory burst that accompanies its absence suggests that *EROS* plays an almost identical role in human and murine immunity. In summary, we have shown that the function of EROS is fully conserved between human and mouse, and that homozygous mutations in EROS underlie a novel sixth cause of chronic granulomatous disease.

## Acknowledgements

D.C.T is funded by a Wellcome-Beit Prize Clinical Research Career Development Fellowship. JCL is a Wellcome Trust Intermediate Fellow. E.C and S.C. are funded by the Wellcome Trust (grant code 098051). A.J.T., GS and AS are supported by both the Wellcome Trust (104807/Z/14/Z) and by the NIHR Biomedical Research Centres of both Cambridge and Great Ormond Street Hospital for Children NHS Foundation Trust/University College London. HAM is funded by the National Science, Technology and Innovation Plan’s (NSTIP) strategic technologies program in Saudi Arabia, (KACST: 13-BIO-755-20). FSA is funded by the Saudi Human Genome Program (King Abdulaziz City for Science and Technology). K.G.C.S. is funded by funded by the Medical Research Council (program grant MR/L019027) and is a Wellcome Investigator and NIHR Senior Investigator.

## Author Contributions

DCT designed and performed experiments and wrote the manuscript. LC analysed patient samples. **A Schejtman** designed guide RNAs, performed CRISPR-mediated deletion of EROS in PLB-985 cells and NBT assays. EC prepared plasmids for transfection experiments and cultured cells. LD performed western blots and assisted with culture of the PLB-985 cells. AB assisted with western blots and ROS assays. JL advised on and performed electroporation of PLB-985 cells. SC and A. Speak advised on cloning strategies and oversaw plasmid preparation. AT and GS oversaw CRISPR-mediated deletion experiments and advised on other experiments. FSA co-ordinated the sequencing of the patient and provided advice on experiments. HA and HAM treated and diagnosed the patient in Saudi Arabia. TC oversaw the clinical care of the patient in Boston and contributed to writing the manuscript. KGCS oversaw experiments in the project and wrote the manuscript.

---

**Figure S1:**
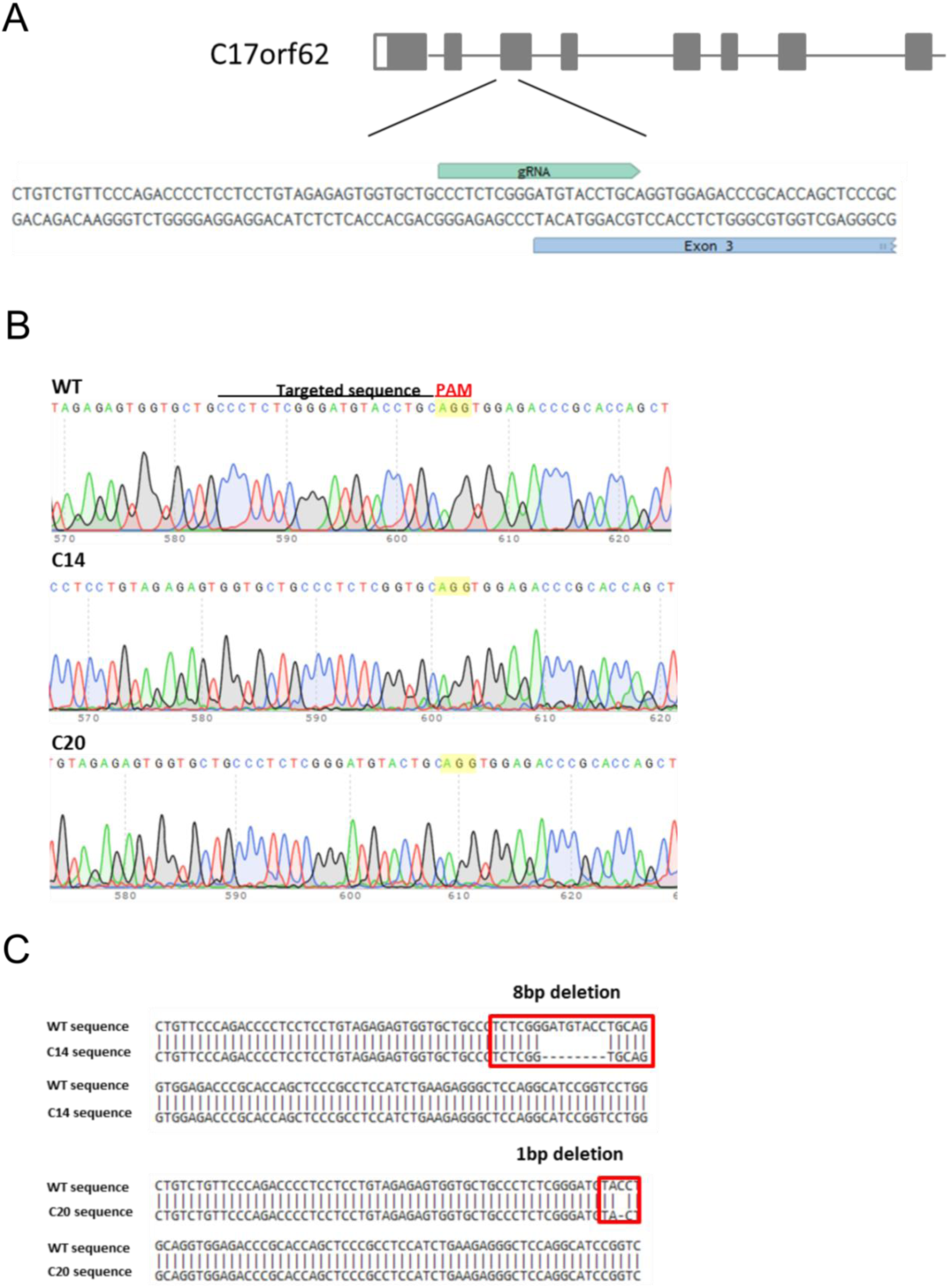
Targeting of EROS for deletion by CRISPR-Cas9. **(A)** Schematic representation of the human *C17ORF62* genomic locus and of the selected guide RNA targeting exon 3 **(B, C)** Sequencing analysis from two clones derived from PLB985 cells that had been electroporated with Cas9:gRNA ribonucleoprotein complex, showing deletion of 8bp in clone 14 and deletion of 1 bp in clone 20.

Abbreviations

CGD: Chronic Granulomatous Disease
EROS: Essential for Reactive Oxygen Species
HLH: haemophagocytic lymphohistiocytosis
NADPH: Nicotinamide adenine dinucleotide phosphate
NBT: Nitro blue tetrazolium chloride
DHR: Dihydrorhodamine
PBMC: Peripheral Blood Mononuclear Cell

